# Cell wall charge gates iron availability in plant roots

**DOI:** 10.64898/2026.03.15.711884

**Authors:** Chenlu Liu, Jedrzej Dobrogojski, Paulina Ramirez Miranda, Dominic Wölki, Marco Marconi, Nicolas Ulbrich, Alberto González-Delgado, Hee Sung Kang, Monika Kubalova, Matyas Fendrych, Berit Ebert, Krzysztof Wabnik, Elke Barbez

**Author notes:** For correspondence: Elke Barbez.

## Abstract

Plants acquire essential mineral nutrients from the soil, yet these elements must first traverse the extracellular matrix of the root before reaching the cell surface. How the physical properties of this extracellular compartment influence nutrient distribution and availability remains poorly understood. In plants, this extracellular matrix is formed by the cell wall, which carries a dynamically regulated negative charge that can change during development and in response to environmental cues. Here we demonstrate that cell wall charge functions as a tunable electrostatic gate that determines how iron is partitioned between retention and bioavailability. This decoupling between iron abundance and availability reveals a fundamental tradeoff imposed by extracellular electrostatics. A mechanistic diffusion–binding model shows that increasing wall charge inherently enhances iron sequestration while limiting its mobility at the cell surface. Genetic perturbation of pectin de-methylesterification validates this principle in vivo. Moreover, iron limitation itself triggers active remodeling of cell wall charge, dynamically shifting the balance toward increased iron accessibility. Together, these findings identify the plant cell wall as an active regulator of nutrient homeostasis rather than a passive barrier. By dynamically modulating extracellular electrostatics, roots control iron partitioning and bioavailability, uncovering a new physical layer of regulation in plant mineral nutrition.

**One-Sentence Summary:** The plant cell wall operates as a tunable electrostatic gate that buffers and releases iron through spatially and environmentally regulated charge dynamics.

## Introduction

The plant cell wall is increasingly recognized as a dynamic extracellular matrix with defined physical and electrostatic properties (Peters et al. 2000, Wolf, 2022; Delmer et al., 2024; Cosgrove, 2024). As the interface between plant cells and the soil environment, it forms a porous and charged scaffold that solutes encounter before reaching the plasma membrane. While transporter-mediated nutrient uptake and intracellular regulation are well characterized, the potential role of the extracellular matrix in shaping ion availability prior to cellular uptake remains largely unexplored. A major determinant of cell wall electrostatics is pectin, a family of acidic polysaccharides that can account for up to 30% of primary cell wall dry mass (Caffall and Mohnen, 2009). Pectins are secreted in a highly methyl-esterified form and subsequently modified by pectin methylesterases (PMEs), exposing negatively charged carboxyl groups (Micheli, 2001). As a result, cell wall charge depends on both pectin abundance and its methyl-esterification status (Stavolone and Lionetti, 2017; Bosch and Hepler, 2005; Giovane et al., 2004).

Given the anionic nature of pectins, we hypothesized that the tunable negative charge of the plant cell wall acts as a biophysical reservoir that modulates extracellular ion availability prior to cellular uptake. Iron provides a particularly relevant test case because its high redox reactivity requires tight spatial control over its distribution (Zahra, 2021; Huang et al., 2024). Using mathematical modeling and genetic perturbation of pectin charge in *Arabidopsis thaliana*, we show that cell wall charge enhances iron retention while reducing the pool of freely available extracellular iron. These findings reveal that the plant cell wall provides an additional layer of biophysical regulation by balancing iron sequestration and availability in the root apoplast.

## Results

### A spatial gradient in cell wall charge aligns with developmental transitions of iron bio-availability

To investigate whether cell wall charge contributes to iron accumulation and distribution along the root, we first assessed its spatial variation in *Arabidopsis*. Col-0 wild-type (WT) seedlings were stained with the fluorescent probe COS^488^, which binds de-methyl-esterified (negatively charged) pectin (Mravec et al., 2014). Confocal imaging revealed a pronounced gradient of cell wall negativity along the root, with highest signal intensities in epidermal cell walls of the meristematic zone that progressively decreased toward the elongation and differentiation zones (Fig. 1A, C). Throughout this study, this readout is referred to as cell wall negativity staining. We next asked whether this spatial gradient in cell wall charge is associated with iron distribution along the root. Spatially resolved ion profiling using laser ablation–inductively coupled plasma mass spectrometry (LA-ICP-MS) revealed that total iron was enriched in the meristematic zone and was significantly lower in the differentiation zone (Fig.1D), closely mirroring the cell wall negativity pattern (Fig. 1A).

**Figure 1.**
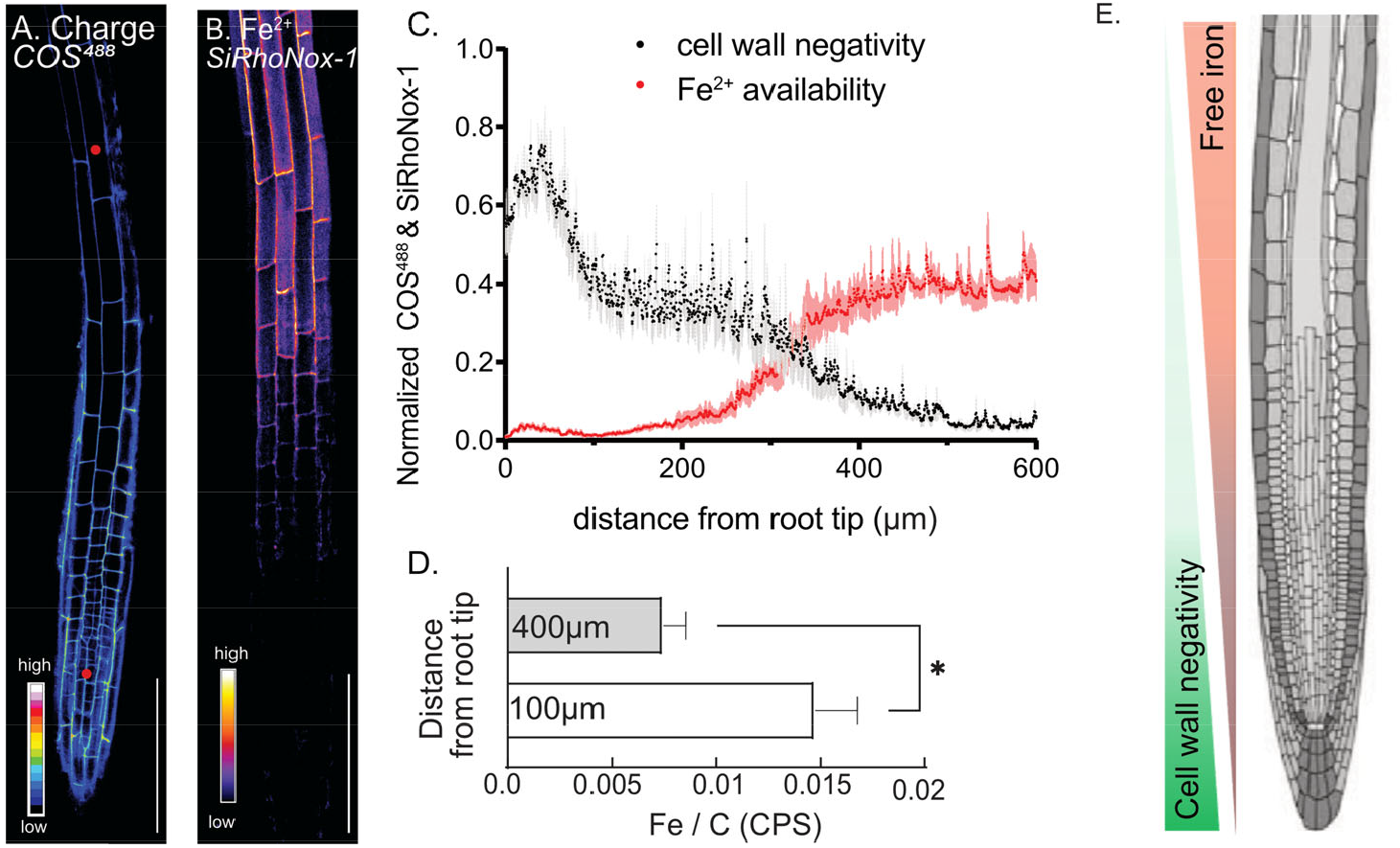
Developmental gradients of cell wall charge correlate with iron partitioning along the root axis. A, Distribution of negative cell wall charge throughout the roots of COS^488^ stained 4-day-old seedlings imaged by confocal microscopy. Color code from black to white indicate low to high COS^488^ signal intensity and thus cell wall negativity. B, Distribution of free Fe^2+^ within the root apoplast throughout the roots of SiRhoNox-1 stained 4-day-old seedlings imaged by confocal microscopy. Color code from purple to white indicates low to high SiRhonox-1 signal intensity and thus free Fe^2+^ levels. C, Quantification of cell wall negativity and Fe^2+^ availability. Black dots indicate the average cell wall negative charge profile from the root tip in shootward direction (error bars indicate s.e.m; n=4 roots); Red dots indicate average free Fe^2+^ profile from the root tip in the shootward direction (error bars indicate s.e.m; n=5 roots). D. Normalized levels of total iron content relative to carbon (counts per second, cps) in the meristematic and differentiation zones of 9-day-old seedlings measured by LA-ICP-MS (error bars indicate s.e.m.; n>5). E, A schematic model illustrating the spatial variation of cell wall negativity and free iron behavior along the root axis during root growth. From root tip to elongation zone, cell wall charge progressively decreases, whereas free Fe^2+^ availability correspondingly increases, indicating an inverse relationship between the two.

Total iron abundance does not necessarily reflect iron availability, as iron can occur in chemically complexed or immobilized states that contribute to total iron levels but are not readily accessible in the extracellular space. To assess the distribution of chemically accessible iron along the root axis, we used the fluorescent probe SiRhoNox-1, which selectively detects reduced Fe^2+^ (Alcon et al., 2024). This readout is referred to hereafter as Fe^2+^ availability staining. In contrast to total iron, Fe^2+^ availability was low in the meristematic zone and increased toward the elongation zone (Fig. 1B, C).

Fe^2+^ availability varied inversely with cell wall negativity along the root axis (Fig. 1A-C). Together with LA-ICP-MS measurements showing elevated total iron levels in regions of high wall charge (Fig. 1D), these results indicate that regions with strongly charged cell walls accumulate more total iron but exhibit lower iron availability, whereas regions with lower wall charge contain more labile iron. These findings reveal a spatial decoupling between total iron accumulation and iron availability along the root axis that correlates with the developmental gradient in cell wall charge. While iron accumulates strongly in the meristematic region, the pool of chemically accessible iron increases toward the elongation zone. Although transporter-mediated processes likely also contribute to iron patterning in roots (Vert et al., 2002), the strong correlation with cell wall charge prompted us to test whether electrostatic retention within the cell wall contributes to this decoupling.

### Charge-dependent interactions in the cell wall create a tradeoff between iron retention and availability

To test whether charge-dependent interactions in the apoplast could explain the decoupling between iron accumulation and iron availability observed above, we asked whether basic biophysical interactions between iron and a negatively charged cell wall matrix could account for this behavior. Pectin—the major anionic component of the primary cell wall—retained iron in vitro, as addition of citrus pectin caused a dose-dependent reduction in soluble iron, indicating electrostatic iron binding (Fig. 2A). We then used these experimentally determined binding properties to parameterize a simplified mathematical model of iron diffusion through a negatively charged matrix (Fig. 2B; see Methods for model details). Briefly, the model represents discrete extracellular compartments (e.g., environment, cell wall, cell interior) and incorporates charge-dependent iron retention within wall compartments, which can generate a locally depleted iron zone in the immediate vicinity of the root. To isolate the specific contribution of cell wall charge, cellular iron uptake was implemented as a set of rate constants that control the abundance of extracellular iron (see Table S2 and Model description in Methods).

**Figure 2.**
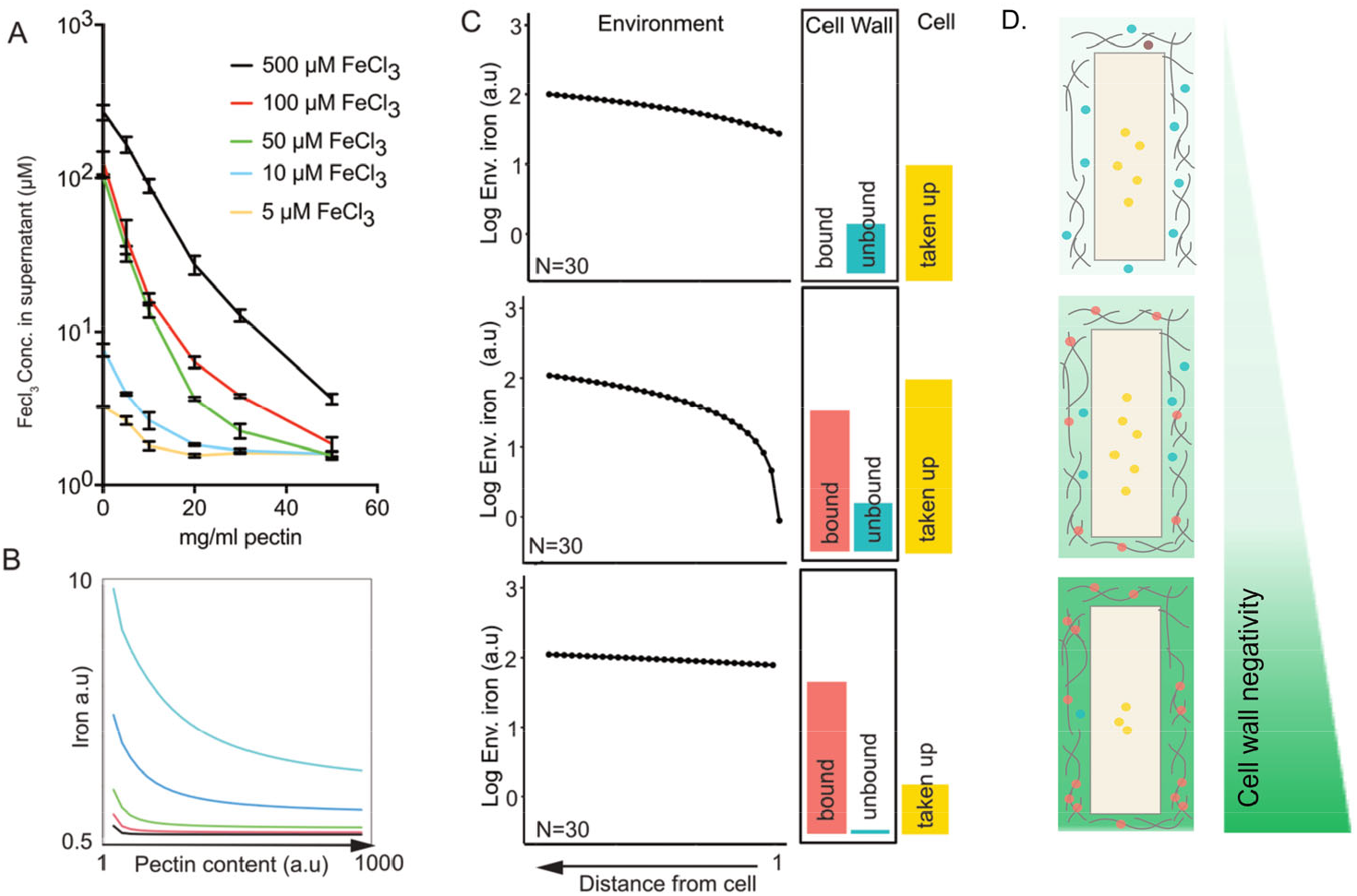
Experimental and modeling analysis of charge-dependent iron retention and mobility. A, Graph depicts the concentration of FeCl_3_ that was not sequestered by pectin in liquid growth medium with the indicated starting concentration of FeCl_3_ and an increasing amount of citrus pectin. (Error bars indicate s.e.m; n=4 technical replicates per condition). B, Predictions of remaining free iron following increased pectin content (cell wall capture) are qualitatively similar to experimental measurements (A). The color of the curves indicates different starting iron concentrations in environment compartments. C, Mathematical model indicating the distribution of generic iron cations in the environment, in the cell wall (bound and unbound) and in the cell in 3 different cell wall charge states: low cell wall negativity (top), medium cell wall negativity (middle), and highly negative cell wall charge (bottom). Error bars indicate s.e.m. D. Cartoon representations illustrate the conceptual interpretation of predicted iron partitioning under different cell wall charge states. Green shading indicates relative cell wall negativity; blue dots represent freely available apoplastic iron, red dots indicate cell wall–associated iron, and yellow dots denote intracellular iron.

Our computational model predicts that increasing cell wall charge enhances total iron accumulation while simultaneously restricting iron availability at the cellular interface. This indicates that an optimal balance between retention and accessibility requires dynamic tuning of wall charge during root growth. Model simulations further revealed a nonlinear dependence of iron partitioning on cell wall charge state. As wall charge increased, the amount of iron retained within wall compartments rose monotonically, reflecting stronger electrostatic association. In contrast, the pool of freely available iron declined despite higher overall iron abundance (Fig. 2C). Together, these results demonstrate that electrostatic retention occurs at the expense of iron availability, establishing a biophysical trade-off between sequestration and accessibility. Based on this principle, we propose a growth-associated capture-and-release mechanism in which highly negatively charged cell walls promote transient iron retention, whereas progressive reduction in wall charge facilitates increased iron availability (Fig. S1).

### Genetic manipulation of cell wall charge uncouples iron accumulation from iron availability in vivo

To experimentally test the model prediction that cell wall charge controls iron retention and availability, we genetically altered pectin-mediated wall charge in Arabidopsis roots. Because pectin de-methylesterification determines the density of negatively charged carboxyl groups, we manipulated the activity of pectin methylesterases (PMEs) and their inhibitors. To increase wall charge, *PME5* and *PME24* were constitutively overexpressed (*PME5ox* and *PME24ox*). To reduce cell wall charge, we used PMEI3 overexpressing seedlings (*PMEI3ox*) and a CRISPR– Cas-generated *pme*^*3,7,24*^ triple loss-of-function line targeting root-expressed PMEs. COS^488^ staining confirmed that PME overexpression increased cell wall negativity epidermal cell walls. *PMEI3ox* and *pme*^*3,7,24*^ roots showed reduced epidermal negative cell wall charge relative to WT (Fig. 3A, B; Fig. S2).

**Figure 3.**
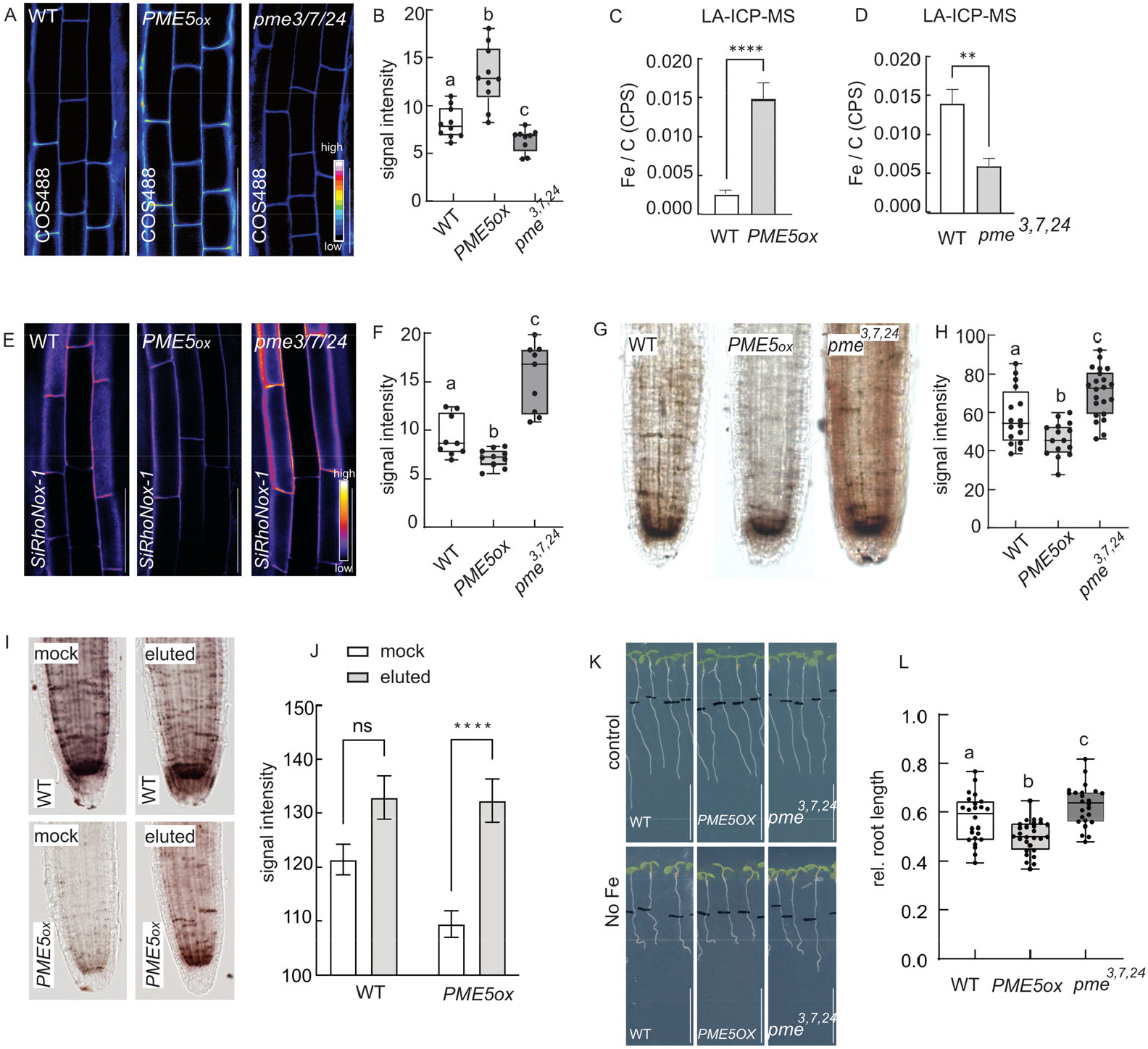
Genetic modulation of cell wall charge alters iron homeostasis and root growth. A, Cell wall charge in roots from 4-day-old WT, *PME5ox* and *pme*^*3,7,24*^ seedlings stained with COS^488^ and imaged via confocal microscopy. Color code from black to white indicate low to high cell wall COS^488^ signal and thus cell wall negativity. B, Box blot indicates the average COS^488^ signal intensity in WT, *PME5ox* and *pme*^*3,7,24*^ roots. (n>9 roots per line). C and D, Normalized levels of total iron content relative to carbon (counts per second, cps) in the meristematic zones of 9-day-old WT, *PME5ox* (C) and *pme*^*3,7,24*^ (D) seedlings as measured by LA-ICP-MS (n>3 roots per line). E, Fe^2+^ availability in cell walls of 4-day-old WT, *PME5ox* and *pme*^*3,7,24*^ stained with SiRhonox-1 and imaged via confocal microscopy. Color code from purple to white indicate low to high SiRhonox-1 signal intensity and thus free Fe^2+^ levels. F, Box blot indicates the average SiRhonox-1 signal intensity in WT, *PME5ox* and *pme*^*3,7,24*^ roots. (n>9 roots per line). G, Perls–DAB staining showing iron distribution in WT, *PME5ox*, and *pme*^*3,7,24*^ seedlings. H, Boxplots show the Perls-DAB staining intensity (n>15 roots per line). I, Perls–DAB staining of WT and *PME5ox* roots under mock conditions (upper panel) or after treatment with 1 mg/mL chitoheptaose for prior to the perls-DAB staining (lower panel). J, Boxplots show the Perls-DAB staining intensity (n>15roots per line). K, L, Root length of 7 day-old WT, *PME5ox* and *pme*^*3,7,24*^ seedlings exposed to iron-deficient growth medium for 3 days relative to seedlings transferred on control (iron-sufficient) medium (n>24). Black lines indicate root tip positions at the time of transfer. Scale bar, 1 cm. Statistical significance was tested using 1-way-ANOVA test, different letters indicate different significant classes (< 0.05) (B, F, H and L), a Student’s t-test (****p<0.0001) (C, D) or a 2-way-ANOVA test with multiple comparison (J). Box limits represent the 25^th^ and 75^th^ percentile, and the horizontal line represents the median. Whiskers display min. to max. values. Error bars represent s.e.m.. Representative experiments are shown. Scale bar, 50 μm (A, E), 1cm (K).

Monosaccharide composition analysis revealed no major differences in overall cell wall polysaccharide content between genotypes (Fig. S3), indicating that these lines primarily differ in wall charge rather than total wall composition. We therefore quantified total iron accumulation using LA-ICP-MS. In line with model predictions, roots with increased wall negativity (*PME* overexpression lines) accumulated significantly more total iron than WT, whereas *pme*^***3***,***7***,***24***^ seedlings showed reduced iron levels (Fig. 3C, D). These data indicate that increased cell wall negativity enhances iron retention within the root apoplast. Because the model predicts that enhanced retention reduces iron availability, we next quantified apoplastic Fe^2+^ using the fluorescent probe SiRhoNox-1. Despite accumulating more total iron, *PME* overexpressors displayed reduced Fe^2+^ availability, whereas seedlings with reduced wall charge (*pme*^***3***,***7***,***24***^ and *PMEI3ox)* showed increased Fe^2+^ levels (Fig. 3E, F; Fig. S4). This data suggests that enhanced wall negativity increases iron accumulation while limiting iron availability.

To assess whether other iron pools are similarly affected, we examined labile Fe^3+^ using Perls–DAB staining. *PME5ox* roots showed reduced staining, whereas *pme*^*3,7,24*^ roots displayed increased signal relative to WT (Fig. 3G, H). Pretreatment with the positively charged oligomer chitoheptaose, which competes with iron for binding to de-methyl-esterified pectin, restored Perls–DAB staining in *PME5ox* roots (Fig. 3I, J), indicating that elevated wall charge reduces iron accessibility through electrostatic retention within the wall matrix.

We next asked whether charge-dependent iron retention has physiological consequences. Root growth was assessed under iron-sufficient and iron-deficient conditions. Under control conditions, all genotypes displayed comparable root growth. In contrast, following transfer to iron-deficient medium, seedlings with highly negative cell walls (*PME5ox* and *PME24ox*) exhibited increased root growth hypersensitivity compared to WT. Conversely, seedlings with reduced wall charge (*PMEI3ox* and *pme*^*3,7,24*^) displayed enhanced root growth under iron deficiency (Fig. 3K, L, Fig. S5A, B, C, D).

Together, these results demonstrate that pectin-mediated cell wall charge directly controls iron retention and availability in vivo and critically determines root growth performance under iron-limiting conditions.

### Iron deficiency triggers pectin turnover and cell wall charge remodeling

Our biophysical findings suggest that modulating cell wall charge can shape iron availability and influence root growth under nutrient stress. If this mechanism is physiologically relevant, plants would be expected to dynamically adjust wall charge in response to changing iron conditions, as proposed by our model (Fig. S1). To test whether plants adapt wall charge in response to iron availability, we exposed seedlings to iron-deficient conditions and analyzed root cell wall negativity. Under iron deficiency, cell wall negativity was reduced, consistent with adaptive remodeling of the cell wall (Fig. 4A, B).

**Figure 4.**
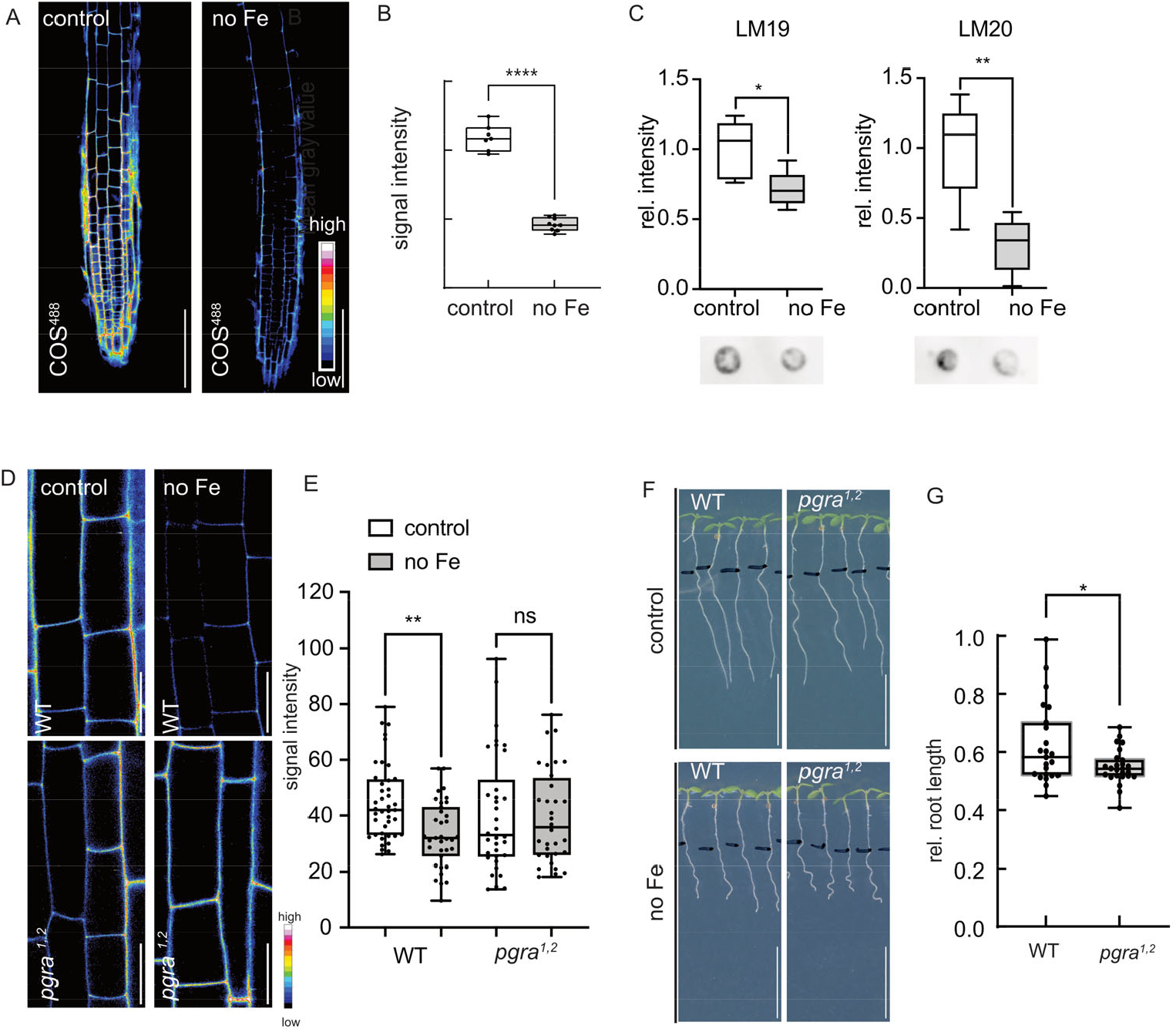
Iron availability dynamically remodels cell wall charge to coordinate root growth. A, Cell wall charge in roots of 4-day-old WT seedlings exposed to iron-deficient medium (no Fe) or control medium for 3 hours. Roots were stained with COS^488^ and imaged via confocal microscopy. scale bar, 100µm. B, Boxplots show the COS^488^ signal intensity in seedlings exposed to control or iron-deficient medium (n>7). C. Dot blot analysis was performed on isolated cell walls of 4-week-old Arabidopsis plants grown in hydroponics exposed to iron deficient medium (no Fe) or control medium for 3 days. Monoclonal antibodies L20 and LM19 were used to detect highly methylesterified (uncharged) and de-methylesterified (charged) pectin epitopes, respectively, n=3. D, Cell wall charge in roots from 4-day-old WT and pgra^1,2^ seedlings exposed to iron deficient medium or control medium for 3 hours and stained with COS^488^ and imaged via confocal microscopy. scale bar, 20µm. E, Boxplots show the COS^488^ signal intensity in seedlings exposed to control or iron deficient medium (n>6). F, G, Root length of 7-day-old WT and pgra^1,2^ seedlings exposed to iron deficient growth medium for 3 days relative to seedlings transferred to control medium (n>13 per line per condition). Statistical significance was tested using Student’s t-test (****p<0.0001) (B, C, G) or a 2-way-ANOVA test with multiple comparisons. (E). Box limits represent the 25th and 75th percentile, and the horizontal line represents the median. Whiskers display min. to max. values. Representative experiments are shown.

A reduction in cell wall charge could in principle reflect altered pectin status under iron deficiency. Hence, we performed dot blot analyses on isolated root cell walls from seedlings exposed to control or iron-deficient conditions using LM19 and LM20 antibodies, which recognize de-methyl-esterified and methyl-esterified pectin, respectively. Roots exposed to iron-deficient medium displayed reduced signals for both LM19 and LM20, indicating an overall decrease in pectin abundance rather than a selective shift in methylesterification state. These findings are consistent with the idea that iron deficiency promotes pectin turnover.

Accordingly, we asked whether the response to iron deficiency depends on pectin degradation. We focused on two root-expressed polygalacturonases, *PGRA1* and *PGRA2*, which degrade de-methyl-esterified pectin (Kubalova et al., 2025). Recent transcriptomic data show that both genes are upregulated under iron deficiency, suggesting that pectin degradation may form part of the adaptive response (Zhang et al., 2025).

Using cell wall negativity staining, we found that the *pgra*^*1,2*^ double mutant failed to reduce wall negativity in response to iron deficiency compared to WT seedlings (Fig. 4D, E). This supports the idea that iron deficiency-dependent degradation of pectin contributes to charge remodeling. Consistent with its functional importance, *pgra*^*1,2*^ seedlings were hypersensitive to iron-deficient conditions relative to WT (Fig. 4F, G). Together, these results identify cell wall negativity as a dynamic property that is remodeled in response to iron availability and contributes to root adaptation under iron limitation.

## Discussion

Here, we identify the plant root extracellular matrix as a biophysical regulator of iron homeostasis that controls iron retention and availability in the root apoplast. This extracellular mechanism complements well-established transporter-mediated and intracellular regulatory systems that coordinate iron uptake and distribution (Bienfait et al., 1985; Giehl et al., 2023; Leskova et al., 2025). Previous studies have shown that iron can associate with cell wall polymers such as hemicelluloses and pectins (Li et al., 2023; Huang et al., 2025). Our findings extend this view by showing that dynamic modulation of cell wall charge actively organizes iron distribution along the root axis rather than passively binding excess metal.

By combining spatially resolved imaging, ion profiling, genetic perturbations, and mathematical modeling, we uncovered a fundamental trade-off between cell wall-mediated iron retention and iron availability. A highly negatively charged wall matrix increases iron retention but simultaneously reduces the pool of iron accessible for cellular uptake. During root growth through the soil, iron encountered by the root tip may therefore become transiently associated with the highly negatively charged cell walls of meristematic cells, effectively creating a temporary extracellular iron reservoir within the apoplast. As these cells leave the meristem and enter the elongation zone, the progressive reduction in wall charge would weaken electrostatic interactions, allowing previously retained iron to become accessible for cellular uptake. In this way, charge-dependent iron partitioning may couple extracellular iron availability to developmental progression along the root axis (Fig. S1).

Consistent with this model, genetic perturbation of cell wall charge confirmed the functional relevance of this mechanism in vivo. Manipulating PME activity or pectate lyase– dependent wall remodeling altered iron accumulation, iron availability, and root growth in a manner consistent with model predictions. Notably, growth phenotypes correlated closely with overall cell wall negativity, regardless of whether wall charge was increased by enhancing de-methyl-esterified pectin or reduced by preventing its removal. These results indicate that the physical charge state of the extracellular matrix, rather than the enzymatic pathway by which it is generated, determines iron availability and root performance under iron-limiting conditions.

## Conclusion

Together, our findings identify the plant cell wall as a dynamic electrostatic interface that regulates iron partitioning in the extracellular space. By modulating the negative charge of pectin polymers, roots can influence whether iron is transiently retained within the apoplast or becomes accessible for cellular uptake. This mechanism introduces an additional layer of iron regulation that operates independently of classical transporter-mediated uptake and instead relies on the physical properties of the extracellular matrix. Because pectin charge is developmentally regulated and responsive to environmental cues, electrostatic control of iron partitioning may provide plants with a flexible strategy to balance iron retention and accessibility during growth and stress. More broadly, these findings highlight the plant cell wall not only as a structural scaffold but also as a biophysical regulator of iron availability at the plant–soil interface.

## Materials and Methods

### Plant material and growth conditions

The coding sequences of *PME5* (*AT5G47500*) and *PME24* (*AT3G10710*) were cloned into the Gateway-compatible binary vector pK7WG2, which drives constitutive expression under the control of the CaMV 35S promoter. The resulting constructs were introduced into the Arabidopsis thaliana Col-0 background via Agrobacterium tumefaciens strain GV3101 using the floral dip method. The *PMEI3ox* line was published previously (Peaucelle et al., 2008). Seeds were surface-sterilized in a sterile laminar flow hood using 70% ethanol and stratified in the dark at 4 °C for 48 hours. Then transferred to a controlled growth chamber maintained at 20 °C under long-day conditions (16 h light/8 h dark) with a light intensity of 100 µmol m^−2^ s^−1^. Seeds were germinated on vertical square plates containing iron-sufficient medium composed of half-strength Murashige and Skoog (MS) salts without FeNaEDTA (Duchefa Biochemie, product no. M0255), 1% (w/v) plant agar (Plant Agar from Duchefa Haarlem, the Netherlands, CAS No. 9002-18-0), 2.5 mM MES, 50 μM ferric sodium EDTA (FeNaEDTA; CAS No. 15708-41-5), and buffer to pH 5.9 using 5 M KOH. The iron-deficiency medium was prepared using the same MS salts, MES, and agar, but 100 μM ferrozine (Sigma-Aldrich, CAS No. 63451-29-6) was added as an iron chelator to bind residual trace iron. Distilled water was used for all media preparations.

CRISPR/Cas9-mediated genome editing was performed using a system adapted from Schiml et al. (2016). Target-specific 20-bp guide sequences were cloned downstream of the AtU6-26 promoter using a Golden Gate strategy. For each gene, two independent gRNAs were designed and assembled into the final binary vector via Multisite Gateway cloning. Constructs were introduced into *Arabidopsis thaliana* (Col-0) by Agrobacterium-mediated transformation. Mutations (insertions/deletions) were confirmed by PCR and Sanger sequencing (Fig. S6). Gene expression levels in mutant lines were validated by RT–qPCR.(Fig. S7)

### Growth assays

Four-day-old Arabidopsis thaliana seedlings grown on iron-sufficient medium were transferred to either control or iron-deficient medium. At the time of transfer, root tip positions were marked. Plates were maintained in a vertical position to allow straight root growth throughout the treatment period. After three days of treatment, plates were scanned using a flatbed scanner, and primary root lengths were measured from the marked position to the root apex using Fiji (ImageJ). Relative root length was calculated for each genotype as the ratio of root length on iron-deficient medium to the average root length on iron-sufficient medium. The experiment was independently repeated three times, with more than 15 seedlings analyzed per line in each repetition. Statistical significance was assessed using one-way ANOVA.

### Confocal microscopy images

Confocal microscopy was performed using a vertical Zeiss LSM 980 Axio Observer Z1/7 system equipped with a Plan-Apochromat 20x and 40×/0.95 Korr M27 air objective. Z-stack images were acquired in line scanning mode to obtain high-resolution optical sections. The excitation and emission settings were as follows: for COS^488^, excitation at 488 nm (laser power 0.7%) and emission collected at 500–550 nm; for SiRhoNox-1, excitation at 633 nm (laser power 12%) and emission collected at 650–700 nm. High-magnification images were acquired at optimized lateral resolution and used for line profile analysis. Seedlings were mounted on agar blocks and imaged with a 1-well glass-bottom chamber. Z-stacks were acquired focusing on the clearest epidermal layer, typically spanning 5–8 optical sections. A subset of 5 optical slices was selected for average intensity projection using Fiji. Each experiment was performed with three biological replicates.

### In vitro pectin binding assays

A series of citrus pectin solutions at concentrations of 0, 20, 40 and 60mg/mL were incubated with a series concentration of FeCl_3_ for 3 hours under continuous agitation. After incubation, samples were centrifuged at 5000 × g for 5 minutes. Then Perls solution (Brumbarova and Ivanov, 2014) was added to the supernatant and incubated for 1 hour. Absorbance at 740 nm was measured using a CLARIOstar microplate reader (BMG LABTECH).

### Plant staining with chemical probes

For COS ^488^ staining, seedlings were incubated in liquid ½ MS medium at room temperature with gentle agitation (50 rpm) for 30 minutes, followed by three washes with the same medium. COS ^488^ was synthesized according to the method described by Mravec (2014). Using a 1:500 dilution from the stock solution. For SiRhoNox-1 staining(Sigma-Aldrich Chemie GmbH, CAS No. 1801528-44-8), the probe was diluted to a final concentration of 5µM in 10mM MES-KOH buffer (pH 6.0). Seedlings were incubated in the staining solution for 3 hours at room temperature in the dark with gentle agitation, followed by three washes in the same buffer.

### Perls-DAB histochemical staining

Perls staining with DAB intensification was performed to visualize labile Fe^3+^ accumulation as previously described (Roschzttardtz et al., 2009) with slight modifications. Briefly, plant tissues were fixed under vacuum in methanol: chloroform: acetic acid (6:3:1, v/v/v) for 1h, and washed 3 times with distilled water. To elute bound iron, tissues were incubated with 1mg ml^-1^ chitoheptaose solution for 1 h followed by incubation in freshly prepared Perls staining solution (4% K_4_Fe(CN)_6_ and 4% HCl, 1:1, v/v) under vacuum at 37 °C for 1 h. After washing 3 times with distilled water, DAB intensification was carried out by sequential incubation in methanol containing 0.3% H_2_O_2_ and 0.01 M NaN_3_ for 1h, followed by incubation in 0.1 M phosphate buffer (pH 7.0) containing 0.025% DAB, 0.005% H_2_O_2_, and 0.005% CoCl_2_ for 3-5 mins. All samples were washed and stored in distilled water prior to imaging.

### Laser ablation ICP-MS

Seedlings were grown on iron-sufficient medium for 7-9 days, then gently washed three times in Milli-Q water. Roots were rapidly frozen in liquid nitrogen and embedded in a epoxy resin. Samples were dried overnight before measurement.

The iron distribution in Arabidopsis thaliana roots were analyzed by LA-ICP-MS, using an Agilent 7900s quadrupole system coupled to an ESI NWR 193 nm laser ablation system at the Department of Geo- and Environmental Sciences (Mineralogy and Petrology) at the Albert Ludwigs Universität Freiburg. Laser ablation spot sizes of 30 μm in diameter were used with a fluence of 0.5 J/cm2 at 7 Hz. The He carrier gas flow was set to 650ml/min. The dwell time was set to 10 ms. A 30 second washout prior and after each analytical period was used to ensure sufficient background levels. SRM NIST 1515 and SRM NIST 610 were analyzed repeatedly throughout the analytical run, with measurements performed after every 15–20 unknown samples to monitor instrumental drift and control signal stability. The instrument was tuned to ThO/Th ratios of <0.3% and U/Th ratios of ∼ 100%, doubly charged cations were tuned to <0.4% using SRM NIST610.

LA-ICP-MS analysis was performed on the root apex and at a region 400–500 µm above the apex.

### RT-qPCR

6-day-old seedlings were harvested in liquid nitrogen and total RNA of dissected roots was extracted using the InnuPREP Plant RNA Kit according to the manufacturer’s instructions [IST Innuscreen, Germany]. First-strand cDNA was synthesized from 1 μg total RNA. Quantitative real-time PCR (RT-qPCR) was performed using SYBR Green chemistry on a CFX384 Touch Real-Time PCR Detection System (Bio-Rad). The thermal cycling conditions were as follows: 95 °C for 2 minutes, followed by 39 cycles of 95 °C for 5 seconds and 60 °C for 30 seconds. Each reaction was carried out in three technical replicates for each of two independent biological replicates. Relative gene expression levels were calculated using the 2^^−ΔΔCt^ method. *EIF4A and UBL5* were used as the internal reference gene. Primer sequences are listed in the Supplementary Table 1.

### Image analysis and quantification

Confocal Z-stacks were processed in FIJI (Schindelin et al., 2012) using average intensity projection. Epidermal cell walls were selected using the segmented line tool, and mean gray values were extracted. Data visualization and statistical analyses were performed using GraphPad Prism 10.

### Cell wall compositional analysis and immunodot blot analyses

Seven-day-old seedlings grown on iron-sufficient medium were transferred to a hydroponic system containing iron-sufficient liquid medium without agar. After 3 weeks of growth, roots were harvested, briefly washed three times with distilled water, immediately frozen in liquid nitrogen, and stored at −80 °C until further use. Roots were subsequently processed to generate alcohol-insoluble residue (AIR) as previously described by Kračun et al. (2017).

For immunodot blot, 2 µl of AIR suspension (1 mg mL^−1^) were spotted onto a nitrocellulose membrane and air-dried for 30 min. Membranes were briefly rinsed with TBS-T and blocked in 5% (w/v) BSA in TBS-T for 30–60 min at room temperature. Membranes were then incubated with LM19 or LM20 (1:100 dilution in 5% BSA/TBS-T) overnight at 4 °C. After three washes with TBS-T (5–10 min each), membranes were incubated with HRP-conjugated anti-rat secondary antibody (1:10,000 dilution) for 30–60 min at room temperature. Following three additional washes with TBS-T, signals were detected using enhanced chemiluminescence (SuperSignal™ West Pico PLUS, Thermo Scientific) according to the manufacturer’s instructions.

For monosaccharide analysis, approximately 2 mg of AIR was divided into six tubes per genotype as technical replicates. AIR pellets were de-starched using 3 U/ml α-amylase, 3 U/ml α-amyloglucosidase and 1 U/ml pullulanase (Megazyme, Cat no. E-BLAAM, E-AMGDF, E-PULBL, respectively), washed once with 96% ethanol, then twice with 70% ethanol and dried overnight (Rautengarten et al., 2019). The de-starched AIR samples were hydrolyzed with 2 N trifluoroacetic acid (TFA) at 120 °C for 1 hour and dried overnight in a vacuum concentrator. The resulting monosaccharides were resuspended in 1 ml of Milli-Q water and analyzed by high-performance anion-exchange chromatography (HPAEC) with pulsed amperometric detection (PAD) on an ICS 5000 system using a CarboPac PA20 (3 × 150 mm) column.

### Mathematical modeling

We developed a mechanistic computational model to simulate the dynamics of cation (i.e. Iron) uptake in a soil-wall-cell interface. The model integrates three interconnected compartments:

1. a spatially explicit soil system,
2. a charged cell wall with dynamic binding properties
3. an intracellular compartment (mimicking cell interior).

The model was implemented in R-studio and simulated using an explicit Euler integrator with a small-time step (*Dt*) of 0.01to preserve stability over a total of 100,000 frames.

#### Numerical implementation

All simulations were implemented in R (version 4.x) using explicit Euler integration with a fixed time step of 0.01. The choice of time step was validated to ensure numerical stability while maintaining computational efficiency. A small numerical constant (ε = 10^−6^) was added when calculating logarithmic quantities to avoid singularities at zero concentrations. Results were visualized using the ggplot2 package, and animations were generated using gganimate to illustrate temporal dynamics.

#### Soil compartment

The soil was represented as a one-dimensional array of (*SoilN*) 30 discrete compartments, each maintaining a dynamic cation concentration (*SoilK*). Cation movement between adjacent compartments is governed by Fickian diffusion with a diffusion coefficient (*SoilD*) of 1.0. For the first soil compartment, we implemented an optional cation production term (*SoilP*) that allowed us to simulate scenarios with continuous cation supply. The diffusion flux between compartments *i* and *j* was calculated as:

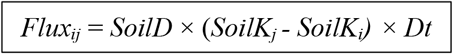

Cation concentrations were constrained to remain between 0 and a maximum value (*SoilMaxK* = 100) to maintain biological realism. The last soil compartment (*SoilN* = 30) interfaced directly with the cell wall, creating a boundary condition for cation transfer.

#### Cell wall compartment

The cell wall was modeled as a single compartment containing two distinct cation pools: bound cations (*WallK*_*b*_) representing cations electrostatically bound to negatively charged pectin molecules, and free cations (*WallK*_*f*_) representing mobile cations in the apoplastic space. The maximum binding capacity of the wall (*WallMaxK*) can be modified across simulations (in the range between 1-1000 units) to approximate the effect of pectin content.

Cation binding dynamics were governed by saturation-dependent absorption and release processes. The absorption rate from soil to bound sites was modeled using a sigmoid function that decreased as the wall approached saturation. The cell wall absorption state follows:

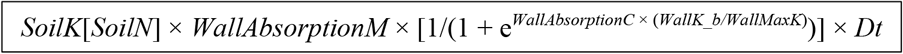

where *WallAbsorptionM* represents the maximum wall absorption rate and *WallAbsorptionC* controls the steepness of the saturation response. Similarly, release from bound to free pools followed a complementary sigmoid function that increased with wall saturation. The release of cation is modelled as:

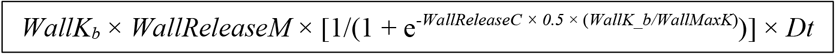

where *WallReleaseM* is the maximum release rate. These sigmoid functions created negative feedback on absorption and positive feedback on release as the wall became saturated, mimicking the finite capacity of binding sites and electrostatic repulsion effects.

The wall also exhibited diffusive exchange with the adjacent soil compartment, with a diffusion coefficient (*WallD*) is adjustable (0.01-1.0) to represent different cation permeability conditions. A binary state variable tracked whether the wall was in “charging” (*WallState* = 0) or “discharging” (*WallState* = 1) mode, switching when bound cation saturation exceeded 99% or fell below 1%, respectively.

#### Cell compartment

The intracellular cation concentration (*CellK*) was governed by uptake from the free wall pool and first-order degradation/efflux processes:

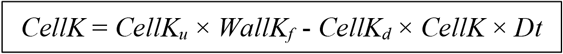

where *CellK*_*u*_ is the cation uptake rate constant and *CellK*_*d*_ is the cation degradation rate constant. The uptake term was proportionally dependent on the availability of free cations in the wall, representing either passive diffusion or facilitated transport processes. Uptake was constrained to never exceed the available free cation pool in any given time step.

#### Membrane potential

The electrical potential difference across the cell membrane (*E*_*m*_) was calculated using a simplified Nernst-like equation based on the concentration gradient between the soil-wall interface and the cell interior:

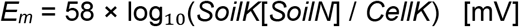

This relationship provided a proxy for the electrochemical driving force for cation uptake, where the factor 58 represents the theoretical slope for a monovalent cation at room temperature (RT/F ≈ 58 mV per decade concentration change).

#### Modelled scenarios

In Fig.2C we present three scenarios for dynamics of cell wall charge and cation uptake by cells and pectin content. First, we systematically varied wall binding capacity (*WallMaxK* from 1 to 1000) in the absence of both soil production (*SoilP* = 0) and cell uptake (*CellK*= 0) to isolate the buffering effect of pectin on residual cation concentrations (Fig. 2B). We then test how cell wall charge and cation viability steer the dynamics of cation uptake and retention (Fig. 2C). Parameter values used to test different scenarios are presented in Table S2.

## Acknowledegements

We thank Jozef Mravec for valuable discussion on cell wall biology and Lothar Kalmbach for the help with the generation of multiplex crispr lines. We thank the LIC Imaging Center Freiburg for expertise and support. We also acknowledge support from the RUB Research School.This project is funded by the Deutsche Forschungsgemeinschaft *(DFG, German Research Foundation):* CIBSS – EXC2189 Project ID 390939984 (to E.B.) and confocal microscopy *Project IDs* 414136422; 499026372. Ministerio de Ciencia Innovación y Universidades of Spain (PID2024-155159NB-I00, CNS2023-143915 to KW), and Severo Ochoa (SO) Program for Centers of Excellence in R&D from the Agencia Estatal de Investigación of Spain [grant CEX2020-000999-S (2022 to 2025) to the CBGP] within the CBGP-CEPLAS International Collaborative Scientific Program grant and Ayudas para contratos predoctorales para la formación de doctores/as 2022 (PRE2022-103239) to AGD. The german research foundation (DFG, 571483527) and the Australian Research Council (ARC, DP220101544) and to BE and HSK.

