## Supplementary information for "Cell wall charge gates iron availability in plant roots"

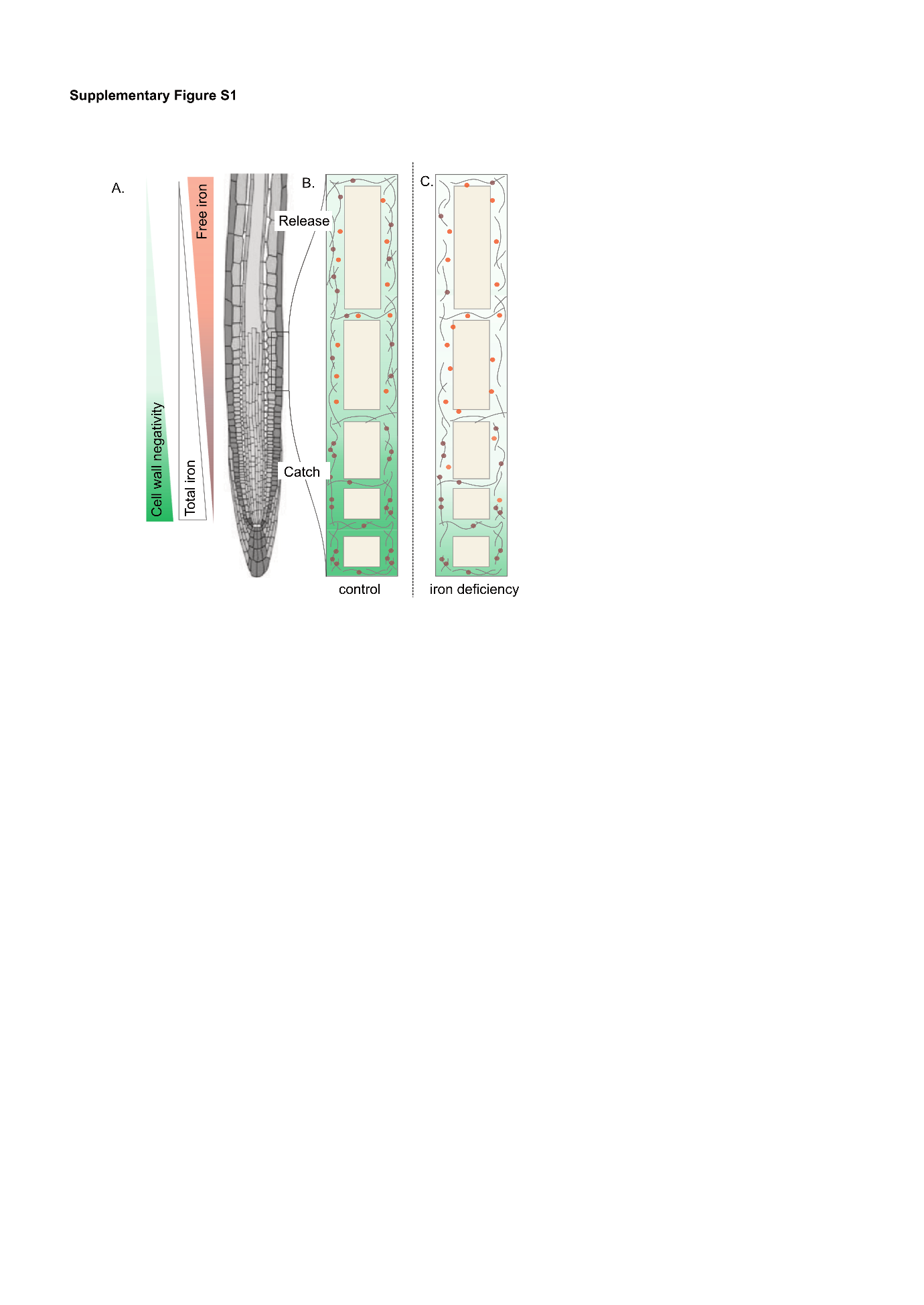

**Supplementary Figure S1. Conceptual model of charge-dependent iron capture and release along the Arabidopsis root axis.**

A, Schematic representation of the Arabidopsis root showing developmental zones and spatial gradients of cell wall negativity, total iron, and free iron along the root axis. Cell wall negativity is highest in the meristematic region and gradually decreases toward the elongation and differentiation zones. Total iron accumulation follows a similar pattern, whereas the pool of chemically accessible iron shows the opposite trend, increasing toward the elongation zone.

B, Conceptual model of charge-dependent iron retention and release under control conditions. In the meristematic region, highly negatively charged pectin polymers in the cell wall promote electrostatic association of iron with the apoplastic matrix, leading to local iron retention. As cells are displaced away from the root tip during growth and enter the elongation zone, cell wall charge progressively decreases. The resulting reduction in electrostatic interactions weakens iron binding to pectin, allowing previously retained iron to be released into the apoplast and become accessible for cellular uptake. This process establishes a developmental capture-and-release mechanism that spatially coordinates iron retention and availability along the root axis.

C, Model of cell wall remodeling under iron-deficient conditions. Iron deficiency triggers pectin turnover and reduces overall cell wall negativity. As a consequence, fewer negatively charged binding sites are available for iron association, resulting in reduced electrostatic retention within the wall matrix. This shift increases the pool of accessible iron in the apoplast and supports iron acquisition under nutrient-limiting conditions.

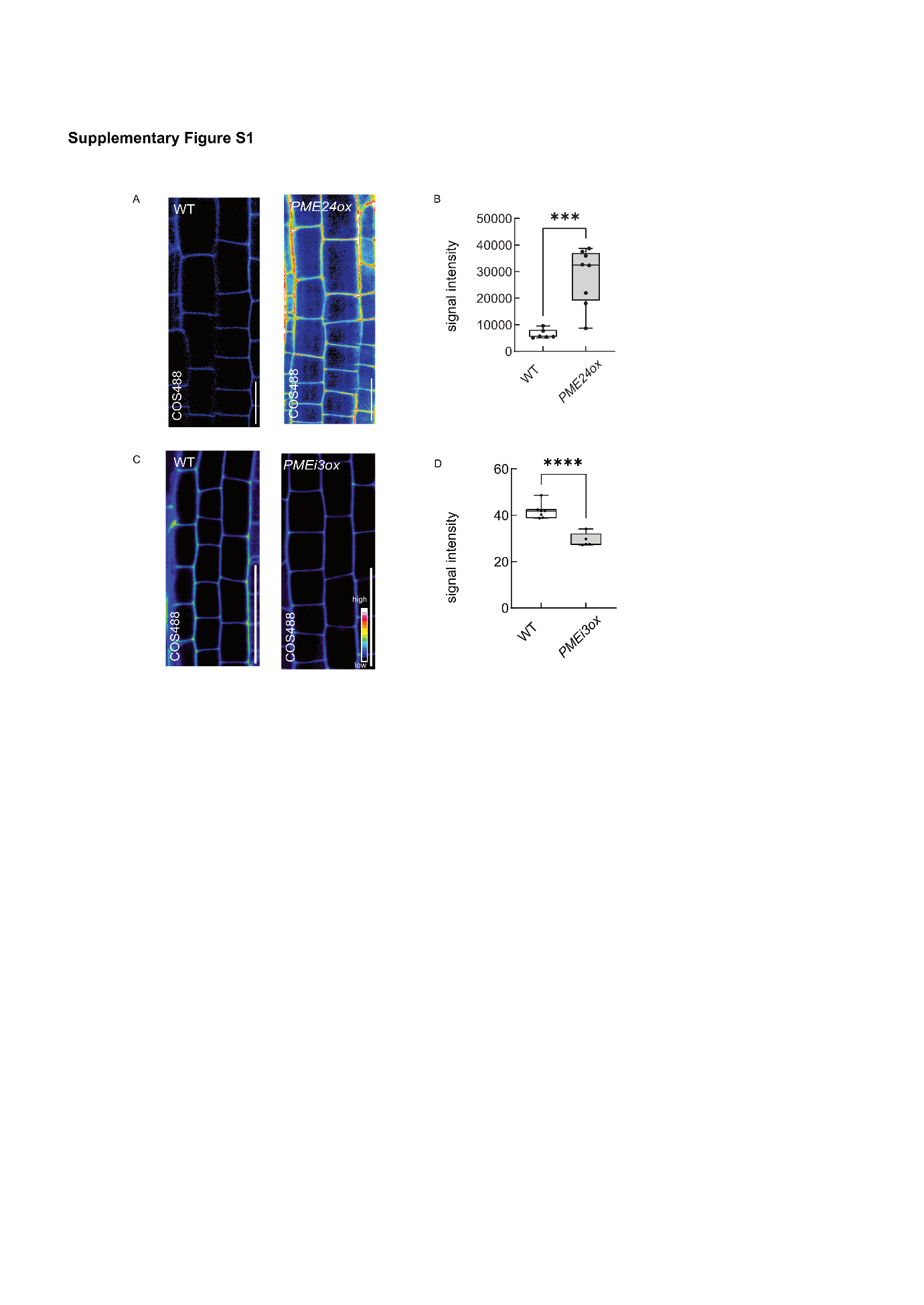

**Supplementary Figure S2, Genetic modulation of PME alter cell wall charge**

Cell wall charge in roots from 4-day-old WT, *PME24ox (A, B)* and *PMEI3ox (C, D)*, seedlings stained with COS^488^ and imaged via confocal microscopy. Color code from black to white indicates low to high cell wall COS^488^ and thus cell wall negativity, scale bar, 20µm. Box blot indicates the average COS^488^ signal intensity in WT, *PME24ox* and *PMEI3ox* roots. (n>7 roots per line). Statistical significance was tested using a Student’s t-test (*****p<0.0001*).

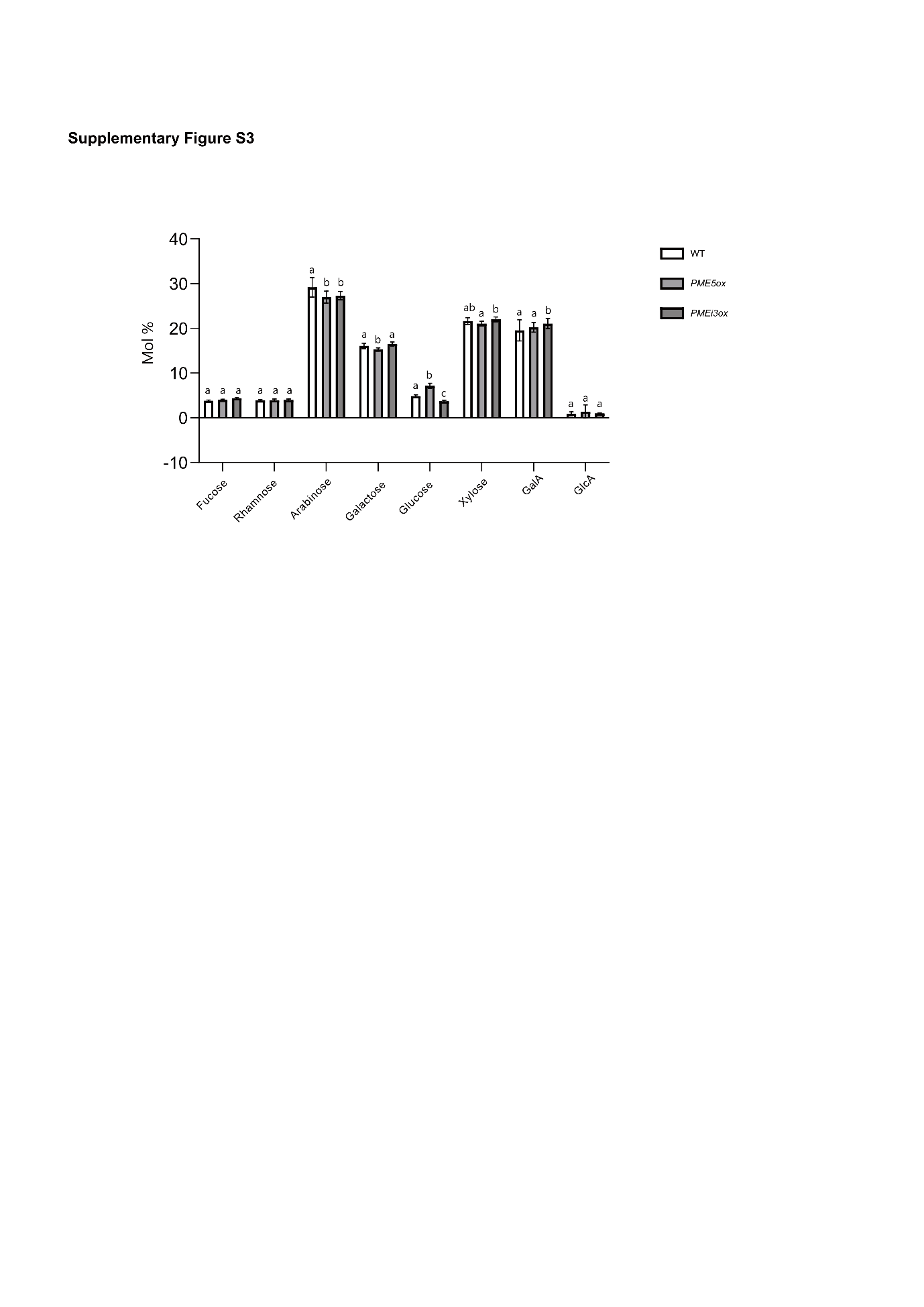

**Fig. S3, Monosaccharide composition analysis of isolated cell walls**

4-week-old Arabidopsis plants grown in hydroponics exposed to iron-deficient medium or control medium for 3 days. Mol% of total monosaccharide content is shown. Statistical analysis was performed using two-way ANOVA followed by Tukey’s multiple comparison test. Different letters indicate statistically significant differences (*P < 0.05*).

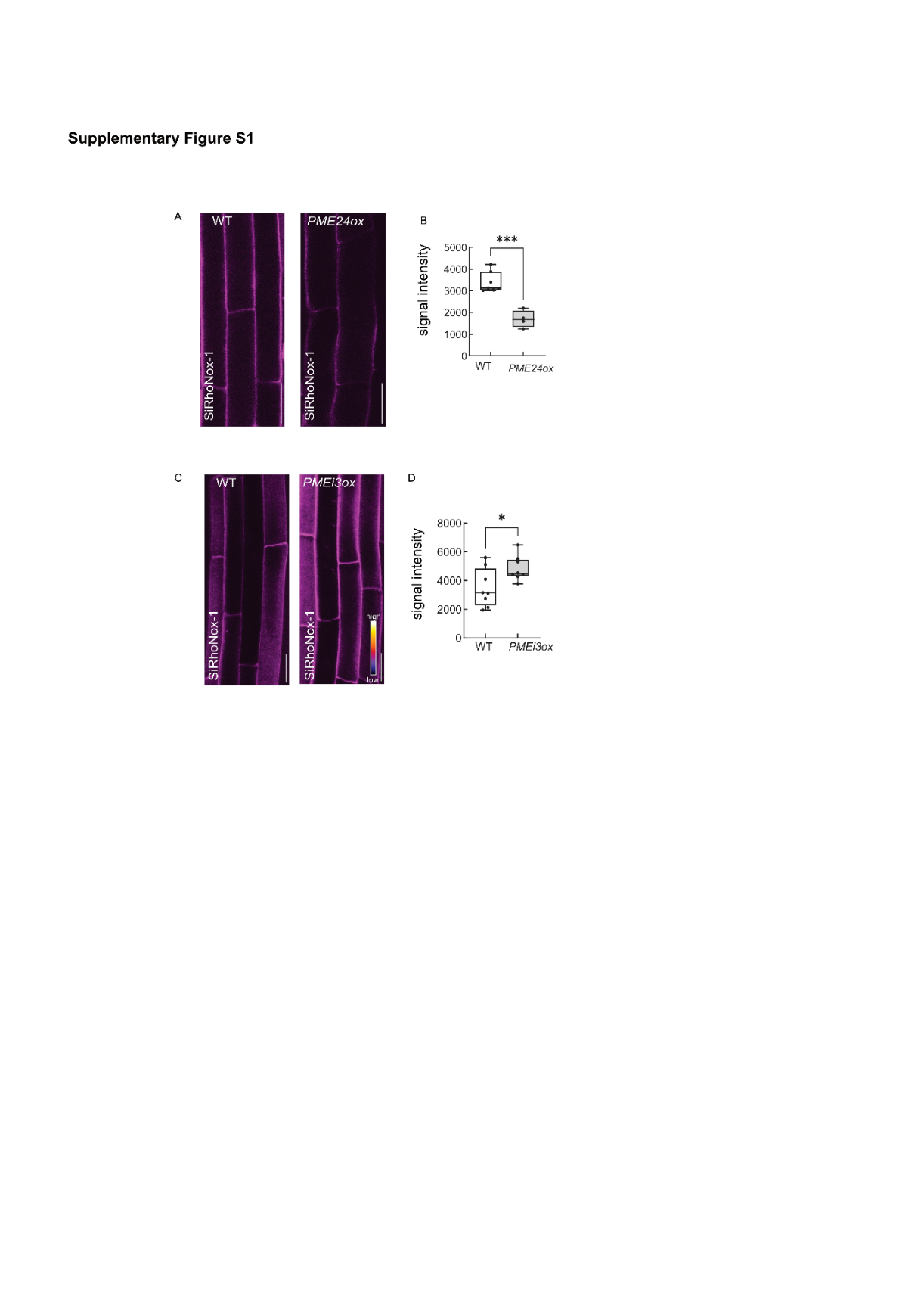

**Supplementary Figure S4, Genetic modulation of PME impact free iron level.**

Fe^2+^ availability in cell walls of 4-day-old WT, *PME24ox (A, B)* and *PMEi3ox (C, D)* stained with SiRhoNox-1 and imaged via confocal microscopy. Color code from purple to white indicates low to high SiRhoNox-1 signal intensity and thus free Fe^2+^ levels. Box blot indicates the average SiRhoNox-1 signal intensity in WT, *PME24ox* and *PMEi3ox* roots. (n>6 roots per line). Statistical significance was tested using a Student’s t-test (****p<0.0001)

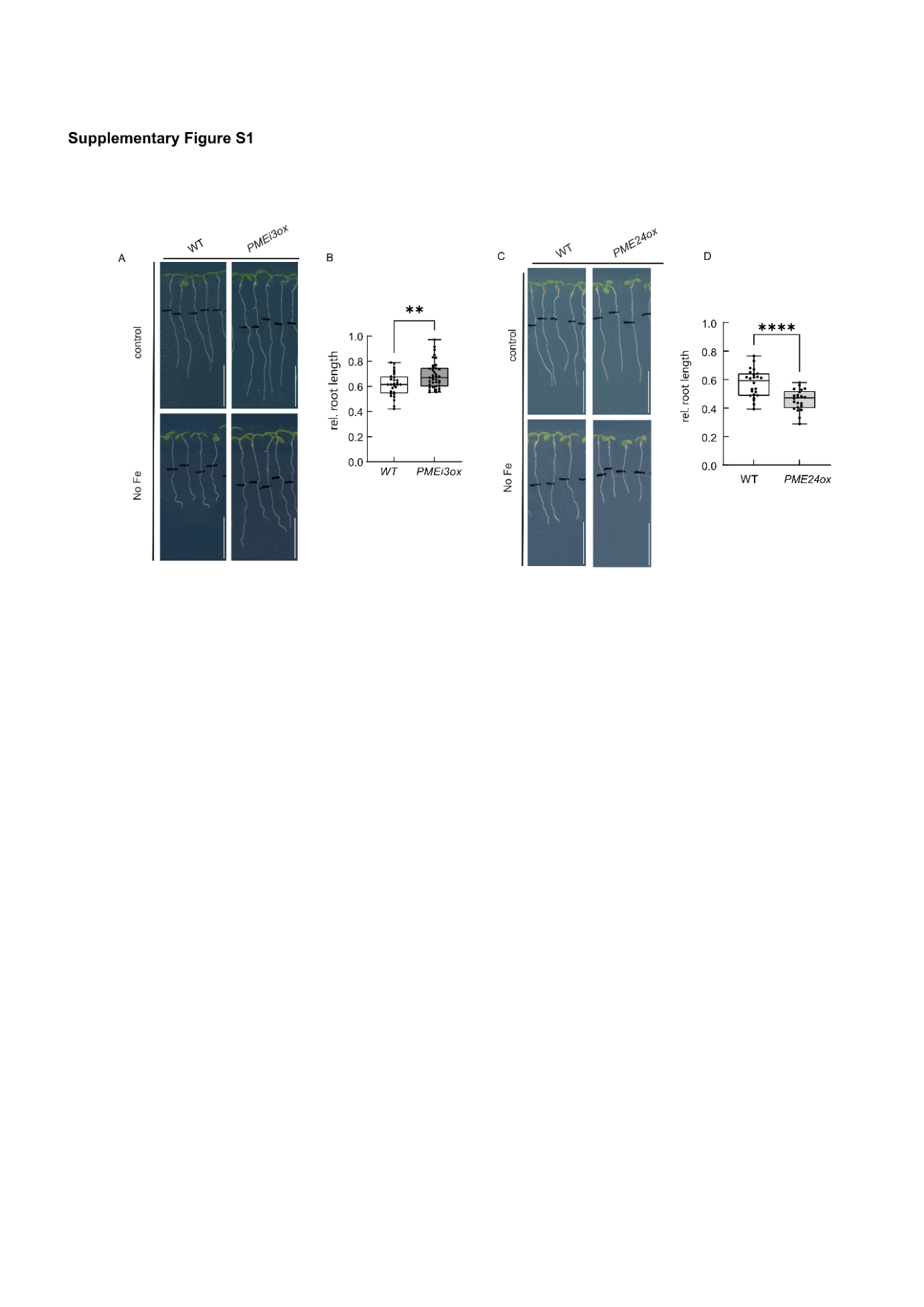

**Supplementary Figure S5, Phenotypic effects of cell wall charge-altered lines under iron deficiency**

Root length of 7 day-old WT, *PME24ox (A, B)* and *PMEI3ox* (C, D) seedlings exposed to iron-deficient growth medium for 3 days relative to seedlings transferred on control (iron-sufficient) medium (n>15). Black lines indicate root tip positions at the time of transfer. Scale bar, 1 cm. Statistical significance was tested using a Student’s t-test (*****p<0.0001*).

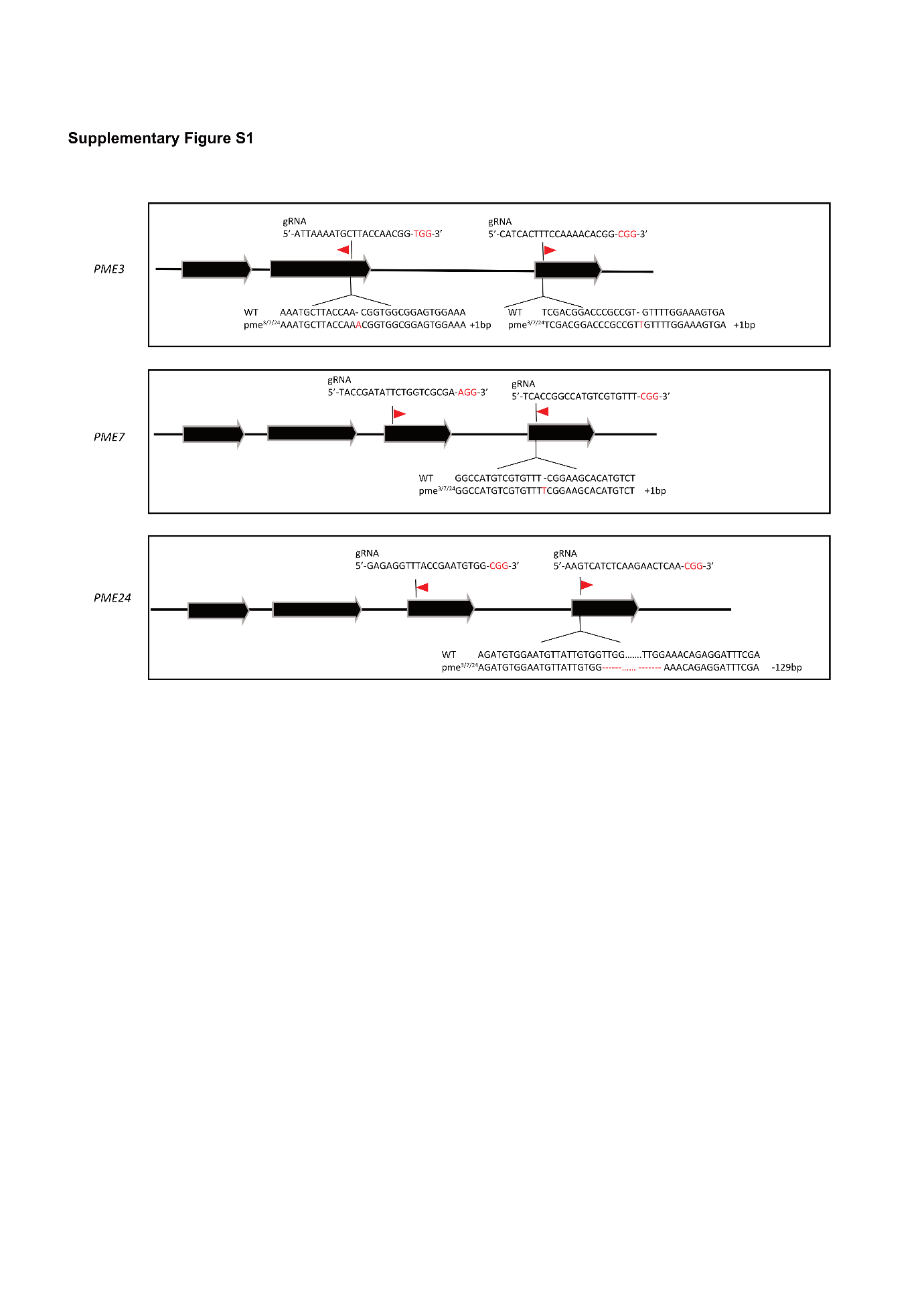

**Supplementary Figure S6, Sequencing confirmation of *pme^3, 7, 24^* and the wild type**

A schematic of the PME loci, CDS region is shown in black arrow, and the mRNA transcripts are indicated by black lines. The positions and sequences of sgRNA target sites are indicated with PAM sequences highlighted in red. Sanger sequencing results for the *pme^3, 7, 24^* triple mutant are shown below the corresponding genomic DNA alignment.

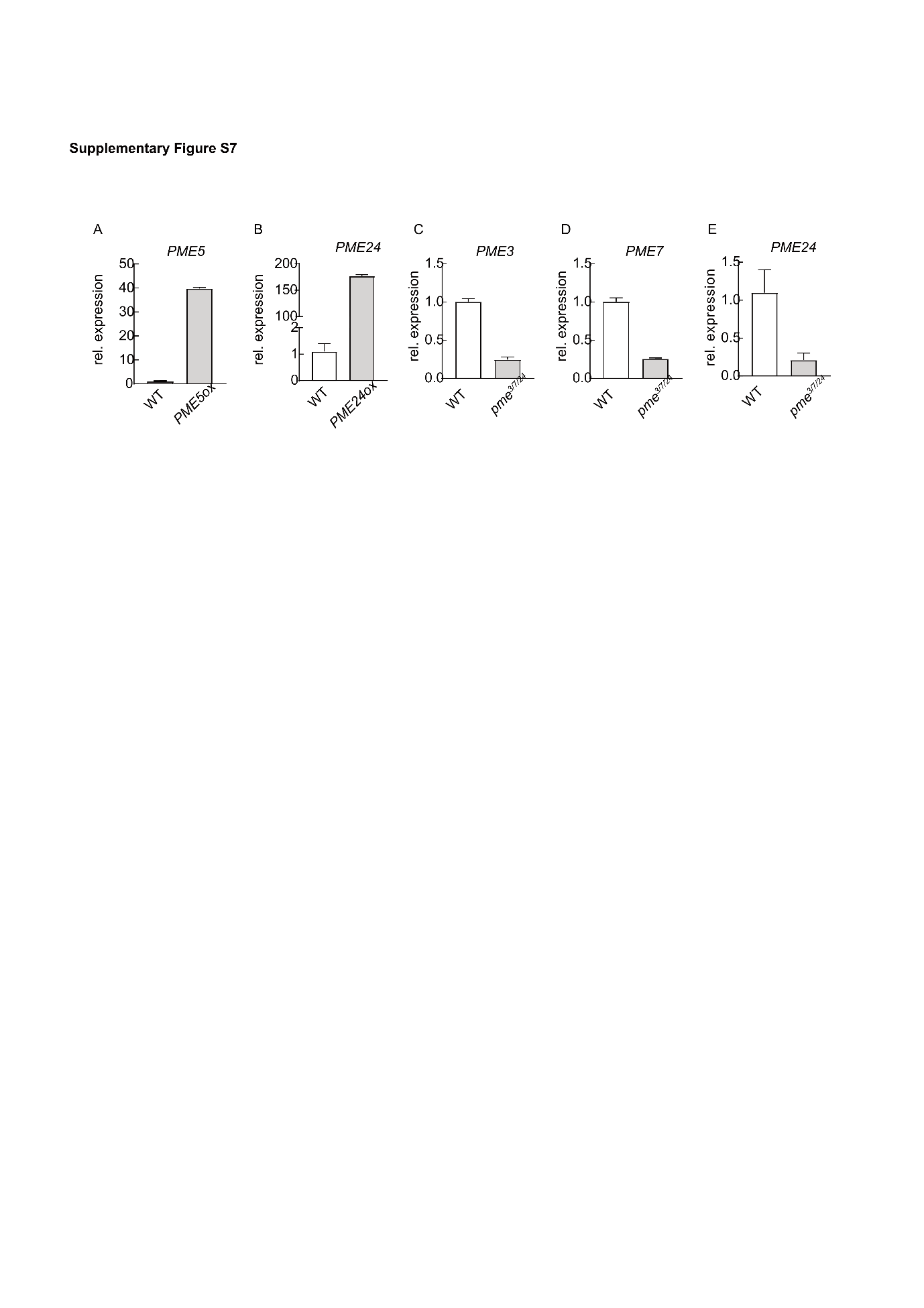

**Supplementary Figure S7, RT–qPCR analysis of PME gene expression in overexpression and CRISPR mutant lines.**

Relative expression levels of *PME5* in *PME5ox* (A), *PME24* in *PME24ox* (B), and *PME3* (C), *PME7* (D), and *PME24* (E) in the *pme3/7/24* mutant were quantified. RNA was extracted from seedlings grown on MS medium for 7 days. *UBL5* was used as the reference gene. Data represent mean ± SE.

**Table S1, Gene and primers used in this study.**

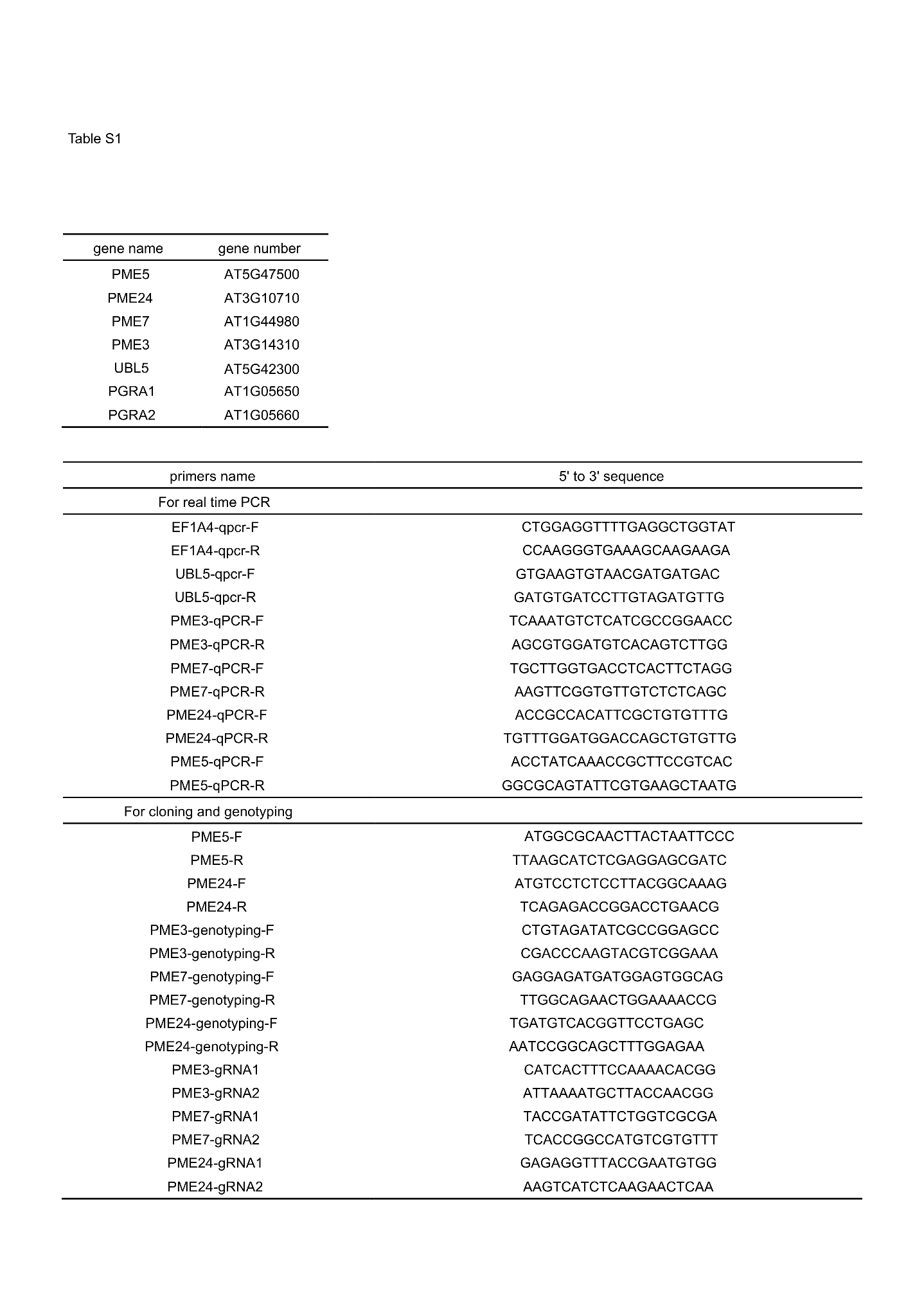

**Table S2, Parameter values used to test different scenarios**

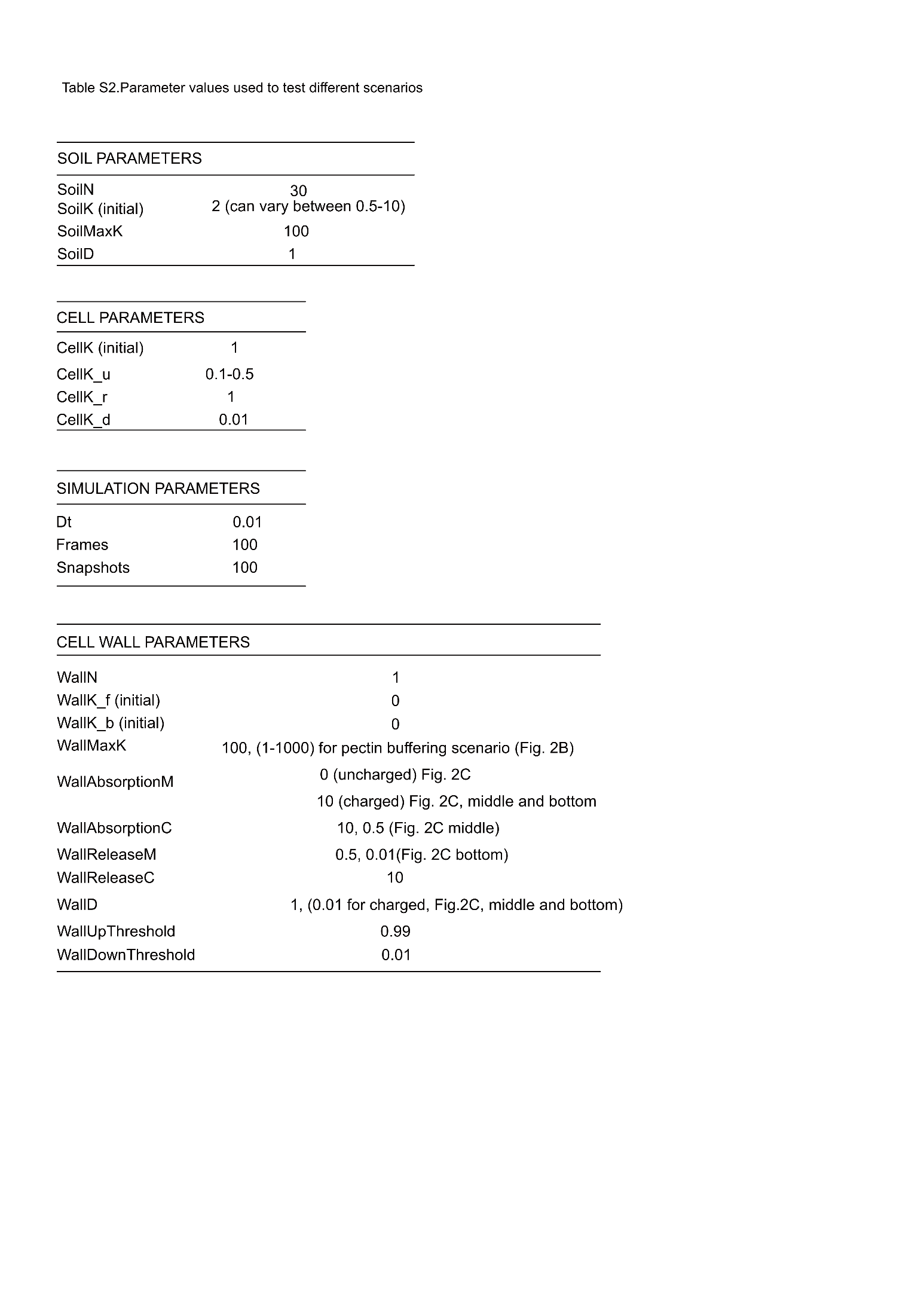
